# Mathematical model of transcription loss due to accumulated DNA damage

**DOI:** 10.1101/2024.07.15.603615

**Authors:** Marko Raseta, Shannon Dealy, Jacinta van de Grint, Jiang Chang, Jan Hoeijmakers, Joris Pothof

## Abstract

We offer a simple mathematical model of gene transcription loss due to accumulated DNA damage in time based on widely agreed biological axioms. Closed form formulae characterizing the distribution of the underlying stochastic processes representing the transcription loss upon specified number of DNA damages are obtained. Moreover, the asymptotic behavior of the stochastic process was analyzed. Finally, the distribution of the first hitting time of transcription loss to specified biologically relevant levels was studied both analytically and computationally on mice data.

## 1 Introduction

Transcription is the process of creating an RNA copy of a gene’s DNA sequence called messenger RNA (mRNA) which represents a carrier of the gene’s protein information encoded in DNA. In humans and other complex biological organisms, mRNA moves from the cell nucleus to the cell cytoplasm, where it is subsequently used in the process of synthesis the encoded protein. DNA damage represents any alteration of the chemical structure of the DNA molecule and can occur both naturally and due to presence of the exogenous factors. Moreover, DNA damages are changes in the structure of the genetic material and can prevent the transcription machinery from functioning and performing properly, causing transcription stress ([24]). Ageing is a naturally occurring biological process associated with a gradual decline in biological function. There is a growing body of scientific evidence that aging is a direct consequence of the accumulation of unrepaired DNA damage. This idea was first suggested in ([1]) with the ever increasing experimental proof over the past decades ([2, 3, 4, 5, 6, 7]). The accumulation of DNA damage is particularly visible in cells that are either non-replicating or slowly replicating, because DNA repair capacity is lower in these cells ([25]). This includes but is not restricted to cells in the brain ([8]), muscle ([9, 10, 11]), liver ([9, 12]) and kidney ([9, 13]). The corresponding reduction in gene expression is observed both on mRNA and protein levels. Further support of the DNA damage theory of aging comes from the observed accelerated aging in humans with inherited defects in DNA repair mechanisms, such as Werner syndrome ([14]), Huchinsosn-Gilford progeria ([15]) and Cockayne syndrome ([16]) with corresponding mean life expectancy of 47, 13 and 13 years, respectively. Moreover, it was recently demonstrated that the age-related transcriptional stress is evolutionary conserved from nematodes to humans. Thus, accumulation of stochastic endogenous DNA damage during aging deteriorates basal transcription, which establishes the age-related transcriptome and causes dysfunction of key aging hallmark pathways, disclosing how DNA damage functionally underlies major aspects of normal aging ([23]).

The purpose of this study is to quantify the connection between the accumulation of unrepaired DNA damage and the associated loss in transcription in a mathematically rigorous fashion and to study the stochastic properties of the associated processes. To be more specific, we quantify the distribution of the transcription loss as a function of the number of accumulated DNA damages in closed form. Furthermore, although the closed form formula for the distribution of the first hitting time of biologically relevant levels of blocked transcription is available, it is impractical due to computational complexity and hence simulation is used to draw inferences on these important distributions. Moreover, we also quantify the distribution of the number of damages needed to switch off both copies of the gene in the genome thereby terminating its biological function.

This manuscript is organized as follows. In Section 2 we list and corroborate the key biological assumptions of the model that are in turn translated into mathematical foundations. Section 3 presents the findings including closed form formulae for the distribution of losses as a function of number of DNA damages accumulated. Section 4 presents findings of a simulation study for the distribution of the first hitting time of the level of accumulated DNA damage to a range of biologically relevant levels. Finally, Section 5 provides some concluding remarks and elaborates on future research directions.

## 2 The set-up

We introduce a mathematical model of transcription loss due to accumulation of DNA damage in biological organisms. DNA damage accumulates at a constant discrete rate ([17, 18]). We shall assume that once certain gene exhibits a damage this damage is indeed permanent and the amount of transcription associated with the gene drops to 0 from that moment onward. There exist two DNA strands, one from each parent while the transcription is only relevant in one direction ([19]). We assume that damage is equally likely to occur on any one of these and hence, at any one time, with probability 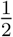, no transcription is lost. Each of the genes will have two copies and the transcription will be equally split between these. The probability that gene *i* will exhibit DNA damage is only assumed to be proportional to its length. Equivalently, we assume DNA is uniformly distributed over the entire length of the genome ([20, 21]).

Consider a biological organism with 2*N* genes in total, thereby taking into account presence of a copy of each gene. For all *i* ∈ {1,…, *N*} let *l*_*i*_ and *α*_*i*_ stand for the length and weight of gene *i*, respectively, with restriction:

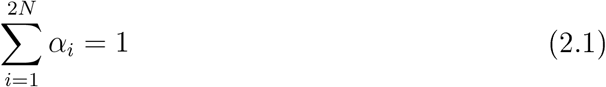

where, for convenience, we rearranged the genome to have *α* _i_ = *α*_*N*+*i*_ for all *i* ∈ {1,…, *N*}. In line with the assumptions above we will assume that, at any instance of time and independently of both past and future, the probability that gene *i* will exhibit DNA damage equals:

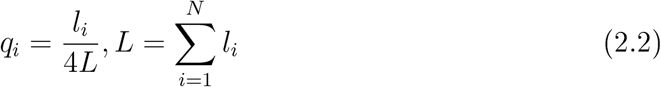

where again *p*_*i*_ *= p*_*N*+i_ for all *i* ∈ {1,…, *N*}.

Let us introduce a partition of the interval [0,1] which we will use throughout the manuscript:

### Definition 2.1.

*For i ∈* {1,…, 2*N* +1} *define the following sequence of intervals*

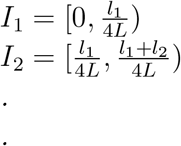

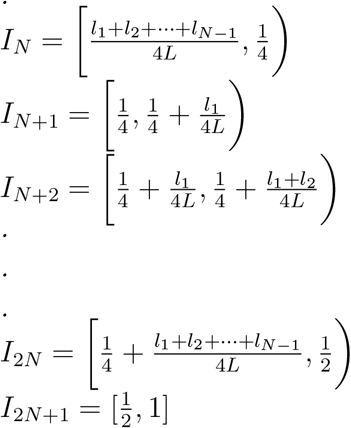

## 3 Results

We begin with a trivial yet important observation.

### Proposition 1.

*Let U be a uniformly distributed random variable on the interval* [0,1]. *Then:*

*(i)* ℙ *(gene i exhibits damage at any time) =* ℙ (*U* ∈ *I*_*i*_)

*(ii)* ℙ (either *one of the two inactive strands is hit at any one time)=* ℙ (*U* ∈ *I*_2*N*+1_

**Proof**. The result follows immediately from the fact that ℙ (*U* ∈ *I*) = |I| for all intervals *I* ⊆ [0,1], where |I| is the length of interval I.

### Definition 3.1.

*Let ω*_*n*_ *stand for the sequence of random variables representing the overall amount of transcription lost after exactly n DNA damages have occurred. Note that if certain gene exhibits more than one damage this has no additional effect as we assume that a single damage is equally harmful and that transcription from that gene is permanently lost*.

### Proposition 2.

*Assume the above set-up. Then the expected transcription lost after n DNA damages reads*

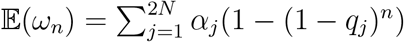

**Proof**. Let 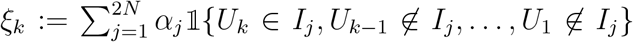, where (*U*_*n*_)_*n*∈ℕ_ is a sequence of independent and identically distributed uniform [0,1] random variables. Observe that Proposition 1 tells us that the entire path of damages can be captured by the sequence (*U*_*n*_)_*n*∈ℕ_. Moreover, we know that the additional transcriptions is lost if and only if a gene which was not previously hit exhibits a damage. Putting these two pieces of information together one can see that *ξ*_*k*_ precisely stands for the amount of additional transcription lost on the *k*^*th*^ DNA damage. Then clearly:

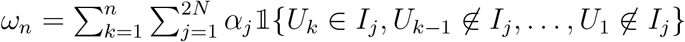

Using the fact that, for all measurable sets, 𝔼 𝟙 (*A*) = ℙ (*A*) we get:

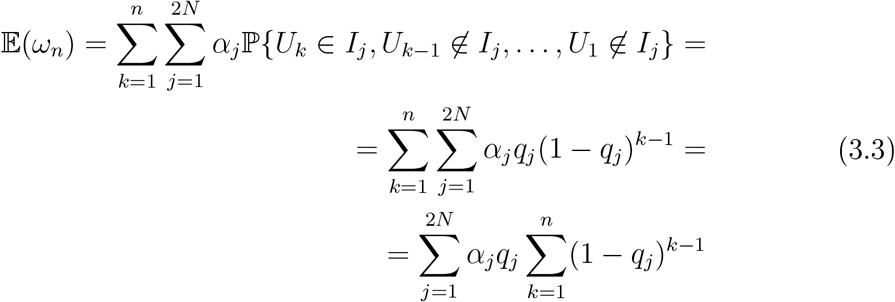

Hence the result follows from the independence of the sequence (*U*_*n*_)_*n*∈ℕ_ and interchanging of the order of summation.

Somewhat more involved calculation yields the behavior of the second moment. More specifically, we have the following result:

### Theorem 3.1.

*There exists a real number τ* ∈ (0,1) *and an absolute constant C such that* 𝕍*ar(ω*_*n*_*) ≤ CT*^*n*^ *for all n simultaneously*.

**Proof**. We begin by computing

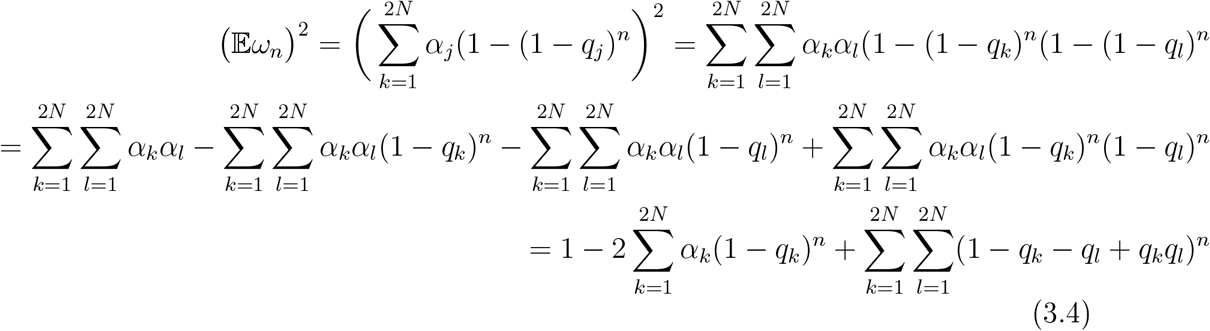

Moreover, we have:

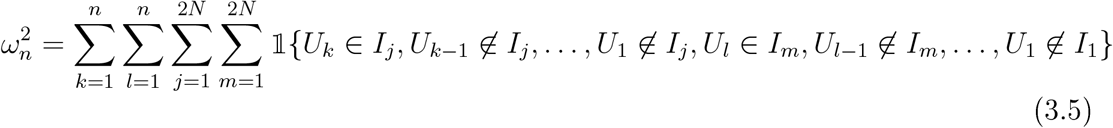

and hence

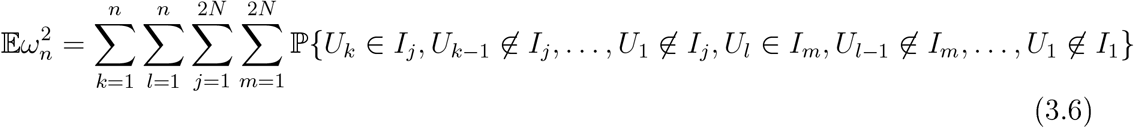

Suppose *k = l* and *j* ≠ *m*. Non-zero contribution of these terms would imply that *U*_*k*_ ∈ *I*_*j*_ and *U*_*k*_ ∉ *I*_*j*_ simultaneously which is clearly impossible. In other words, if *k =* l then only the only cross terms which contribute are those when *j* = *m* as well. We split the sum repeated sum above into three sub-cases, namely *k = l, k > l* and *k < l*. We shall deal with the first two in detail while the third one is easily computed by symmetry. To this end we have:

Case 1: If *k =* l then *j* = m and whence by using the independence and distributional equality of the *Uj*’s together with interchanging the order of summation, we see that the corresponding cross terms reduce to:

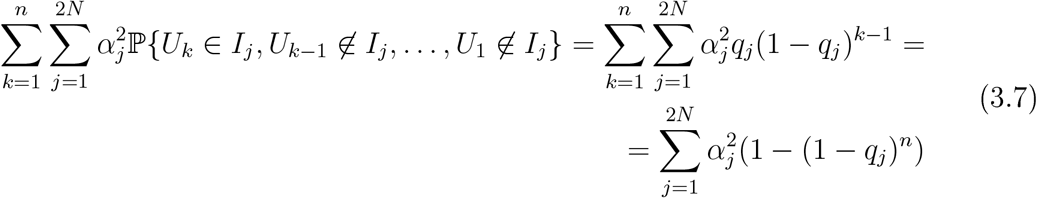

Case 2: *k > l*. The corresponding cross terms read:

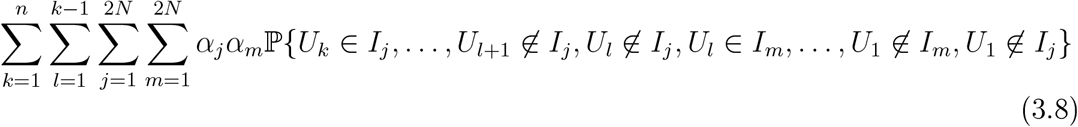

Notice that if *j* = *m* implies that *U*_*l*_ ∉ *I*_*j*_ and *U*_*l*_ *∈ I*_*j*_ and thus those terms with *j* ≠ *m* will yield zero contribution. Moreover, observe that {*U*_*l*_ ∈*I*_*m*_*}* implies that {*U*_*l*_ ∉ *I*_*j*_} since these intervals are disjoint by the very construction. In other words, {*U*_*l*_ ∈ *I*_*m*_*}* ⊆ {*U*_*l*_ ∉ *I*_*j*_} for all *j* ≠ m. This implies that (1.8) simplifies to:

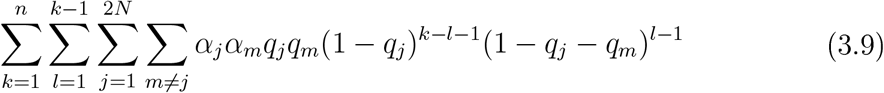

Furthermore, simple algebra and interchange of the order of summation also yields:

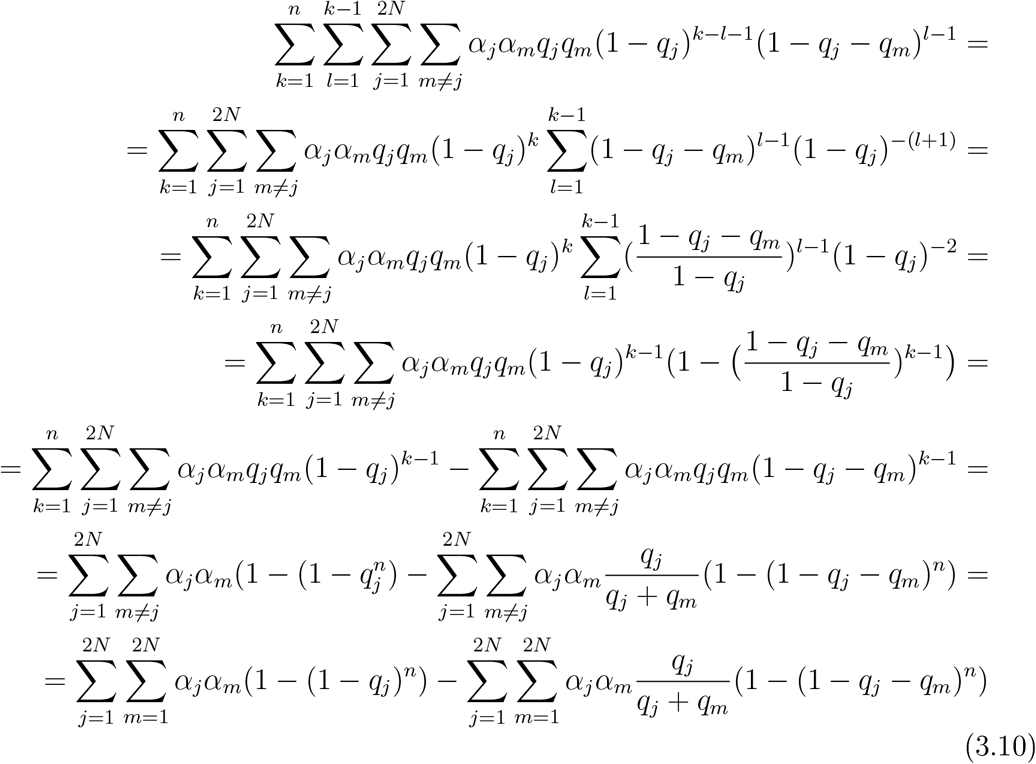

Finally, we shall put this expression in the form when the summations index *j* is unrestricted and hence the expression (3.10) further simplifies to:

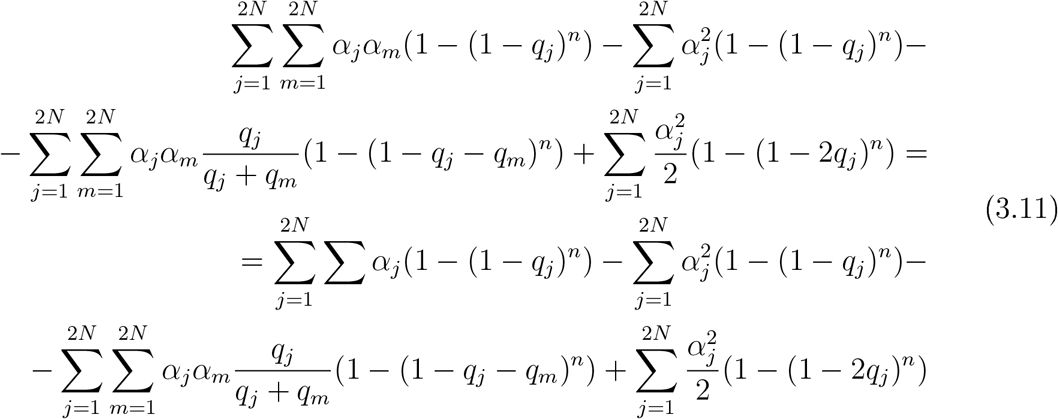

By symmetry, the cross-terms corresponding to the those indices such that *l* ≥ *k* read:

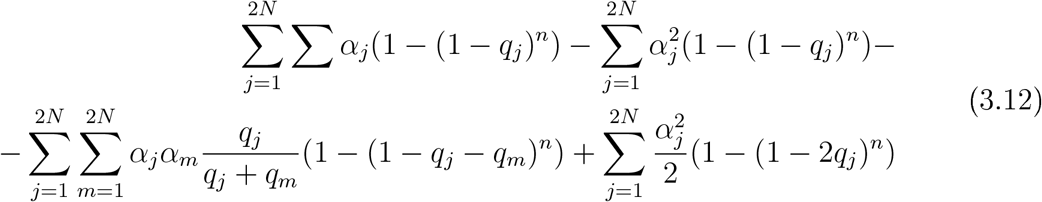

Using suffix notation and summation convention we finalize the expression for the second moment of *ω*_*n*_. Indeed, we have:

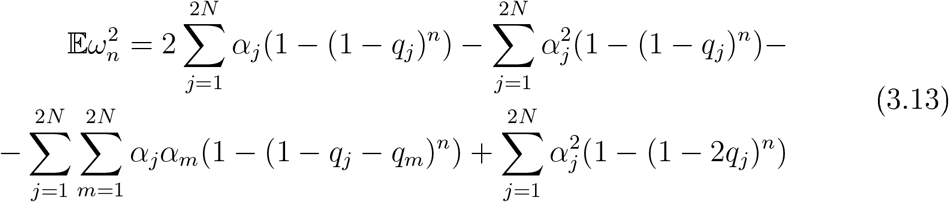

Further simple algebra yields the expression for the variance of *ω*_*n*_. Indeed, we have:

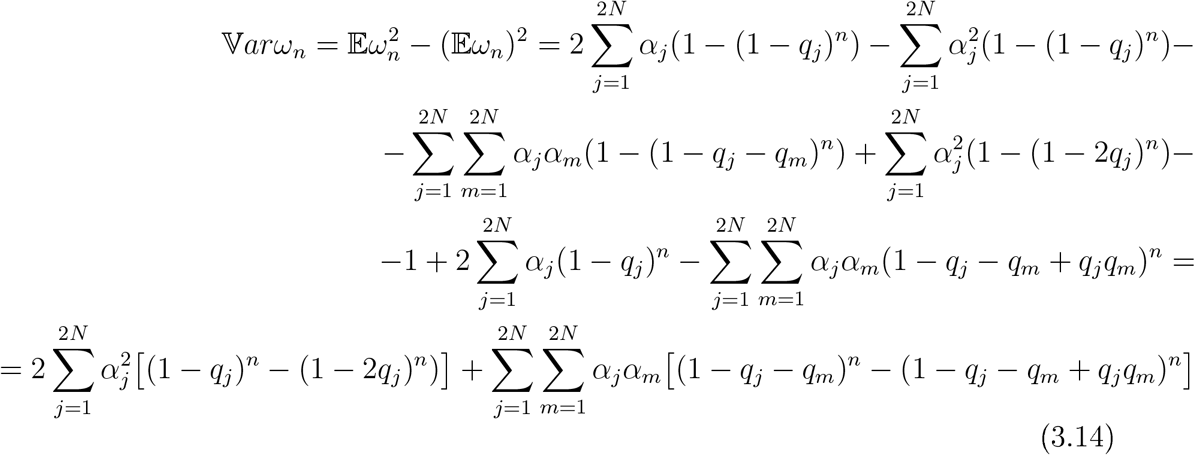

Moreover let us define:

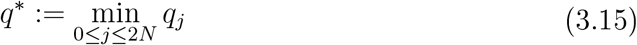

Several applications of triangle inequality finally yield:

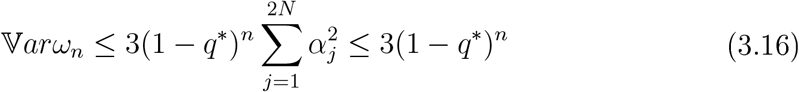

and hence the proof is complete.

As damages accumulate the overall amount of transcription lost increases and eventually approaches 1 (100%). We have seen that lim_*n*→∞_ 𝕍*ar*(*ω*_*n*_) = 0 and lim_*n*→∞_ 𝔼 (*ω*_*n*_) = 1 and hence the model is in line with these simplistic demands. However, much more is true but further definitions and results are needed. To this end we have:

### Definition 3.2.

*A sequence of random variables (X*_*n*_)_*n* ∈*ℕ*_ *is said to converge in L*_*p*_ *to some random variable X, written* 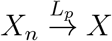, *if and only if* lim_*n*→∞_ 𝔼 (|*X*_*n*_ — *X* |^*p*^ = 0

### Definition 3.3.

*A sequence of L1 random variables (X*_*n*_)_*n* ∈*ℕ*_ *is said to be uniformly integrable if and only if*

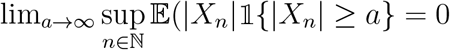

### Definition 3.4.

*We say that a sequence of random variables (X*_*n*_)_*n* ∈ℕ_ *converges to a random variable X in probability, written* 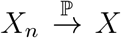 *if and only if for all* ϵ > 0, lim _*n*→∞_. ℙ (|*X*_*n*_ — *X*| ≥ ϵ) = 0

### Definition 3.5.

*A sequence of random variables (X*_*n*_)_*n* ∈ℕ_*converges almost surely to a random variable X, written* 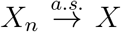 *if and only if there exists some measurable set A with* ℙ (*A*) = 1 *and X*_*n*_*(w) — X*(*w*) *for all w* ∈ *A*.

### Proposition 3.

*Let* (*X*_*n*_)_*n*∈ ℕ_ *be a sequence of random variables and suppose there exists a dominating L1 random variable Y with* |*X*_*n*_| ≤ *Y for all n* ∈ ℕ. *Then the family (X*_*n*_)_*n* ∈ℕ_ *is uniformly integrable*.

**Proof**. Please see page 183 of ([22]) for details.

### Proposition 4.

*Let (X*_*n*_)_*n* ∈ℕ_ *be a monotone sequence of random variables that converges in probability to some random variable X. Then* (*(X*_*n*_)_*n* ∈ℕ_ *converges to X almost surely*.

**Proof**. Please see page 195 of ([22]) for details.

### Theorem 3.2.

*Suppose p* ≥ 1 *and (X*_*n*_)_*n* ∈ℕ_ ∈ *L*^*p*^. *The following statements are equivalent*

*(a) (X*_*n*_)_*n* ∈ℕ_ *is L*_*p*_ *convergent*.

*(b)* 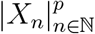 uni*formly integrable and (X*_*n*_)_*n* ∈ℕ_ *converges in probability*.

**Proof**. Please see page 194 of ([22]) for details.

### Theorem 3.3.

*The following statements hold true:*

*(a)* 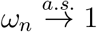

*(b)* 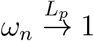

**Proof**.

By definition *w*_*lt*_ ≤ 1 and whence

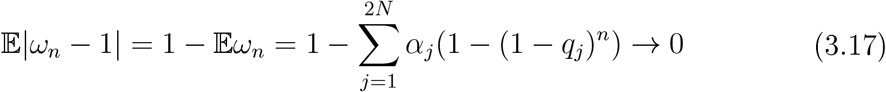

as *n* — ∞ since all genes have strictly positive lengths. In other words, 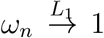. Then by Theorem 1.2 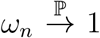. However, as damages can only accumulate in time, *w*_*n*_ ≥ *w*_*n*−1_ for all *n* ∈ ℕ and hence (*ω*_*n*_)_*n*∈ ℕ_ is a monotonically increasing. Thus, by Proposition 4, 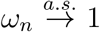. Moreover, as (|*w*_*n*_| ≤ 1 we have (*ω*_*n*_)_*n*∈ ℕ_ ∈ *L*^*p*^ for all *p* ∈ ℝ+. Moreover by Theorem 1.2 (|(|*w*_*n*_|^*p*^)_*n*∈ ℕ_ is uniformly integrable and the proof is complete.

We are also interested in studying the least amount of damages needed to block some prescribed level of transcription,, β, say. More specifically, the levels *β* ∈ {0.3,0.5, 0.7, 0.9} are of biological interest. Mathematically, this is captured by studying the random variable *T* which is the first hitting time of the sequence (*ω*_*n*_)_*n*∈ ℕ_ to level *β*-More specifically:

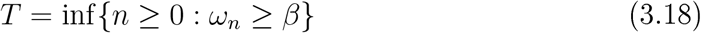

We now provide a closed form expression for the distribution of *T*. By using the law of total probability we have:

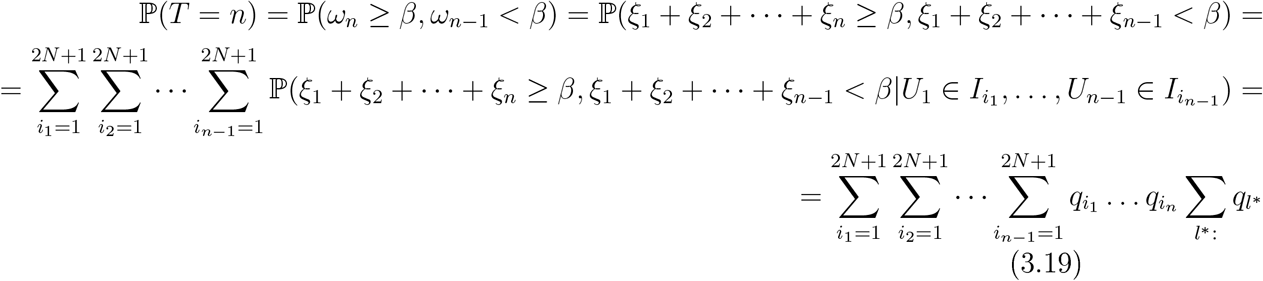

where *l\** corresponds to all those indices satisfying the following relation:

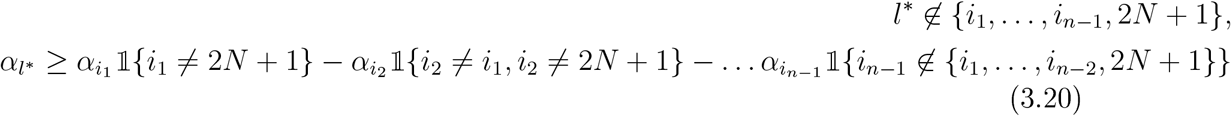

We are also interested in a problem of computing the probability distribution of the number of damages needed to ‘‘switch off” both copies of some specific gene, that is number of damages needed to see the transcription associated with this gene falling to 0. To be more specific, we have:

### Proposition 5.

*Let* Ω* *stand for the random variable representing the smallest number of damages needed for both copies of gene i (in further text these will be labeled i and i*) to exhibit a damage, that is, the first moment in time when the transcription associated with this gene falls permanently to 0. Then:*

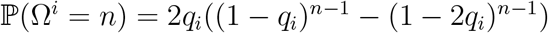

**Proof**.

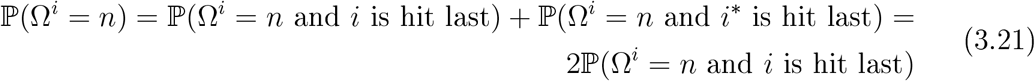

as *i* and *i*\* are equally likely to exhibit a DNA damage. Observe that the {Ω* — n and *i* is hit last} can only occur if and only if among the first *n* — 1 damages at least one occurs in gene *i*\* while gene *i* is hit for the first time precisely on the *n*^*th*^ DNA damage. By conditioning on the number of *i*\**s* among the first *n* — 1 DNA damages we have:

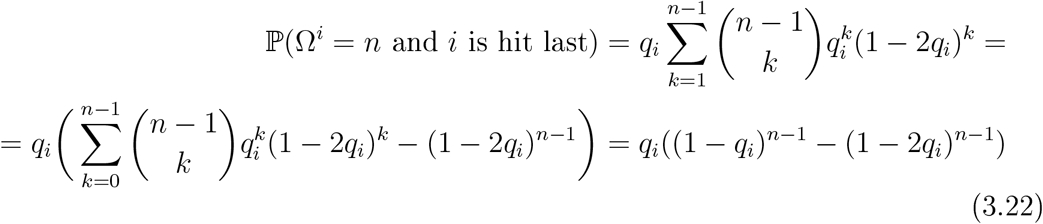

and the proof is complete.

## 4 Simulation study

Although the exact expression for the probability distribution of the random variable *T* is available, its practical value is rather questionable due to the apparent computational complexity needed to implement the closed form solution. However, we provide a simple algorithm which yields the approximate distribution of *T* in a computationally feasible manner. Indeed, we simulate the process (*ω*_*n*_)_*n*∈ ℕ_ specified number of times and record the value of *T* on each such run to obtain the corresponding histograms. The pseudo-algorithm is presented below:

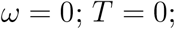

Generate a uniform [0,1] random variable *U*_1_ Find *n* such that *U*_1_ ∈ *I*_*n*_ for *n ∈*

{1,2,…, *2N* + 1}

If *n* = 2*N* + 1 T + +;

else *ω* = *ω* + *α*_*n*_

If *ω* ≥ 5, Stop and return *T* = 1 else generate a random variable *U*_*2*_, *U*_*2*_ independent of *U*_1_ and uniformly distributed on [0,1]

Find *m* such that *U*_*2*_ ∈ *I*_*m*_

if (*m* = 2*N* + 1 or *U*_*1*_ ∈ *I*_*m*_*) T++* ;

else *ω* = *ω* + *α*_*m*_

if *ω* ≥ *β* T=2;

otherwise continue generating new independent and identically distributed uniform [0,1] random variables until you eventually reach the prescribed level *β* return T;

Transcription loss is computed based on relative levels of nascent RNA transcription for each gene from three biological replicates. Data set spans 1331939K total base pairs, 9661 Genes, including both alleles of each gene. The instances of DNA damage were inflicted uniformly at random throughout the genome until a specified loss of transcription has been reached. This procedure was repeated 100 times to generate a representative histogram of the first hitting time *T*.

**Figure 1.**
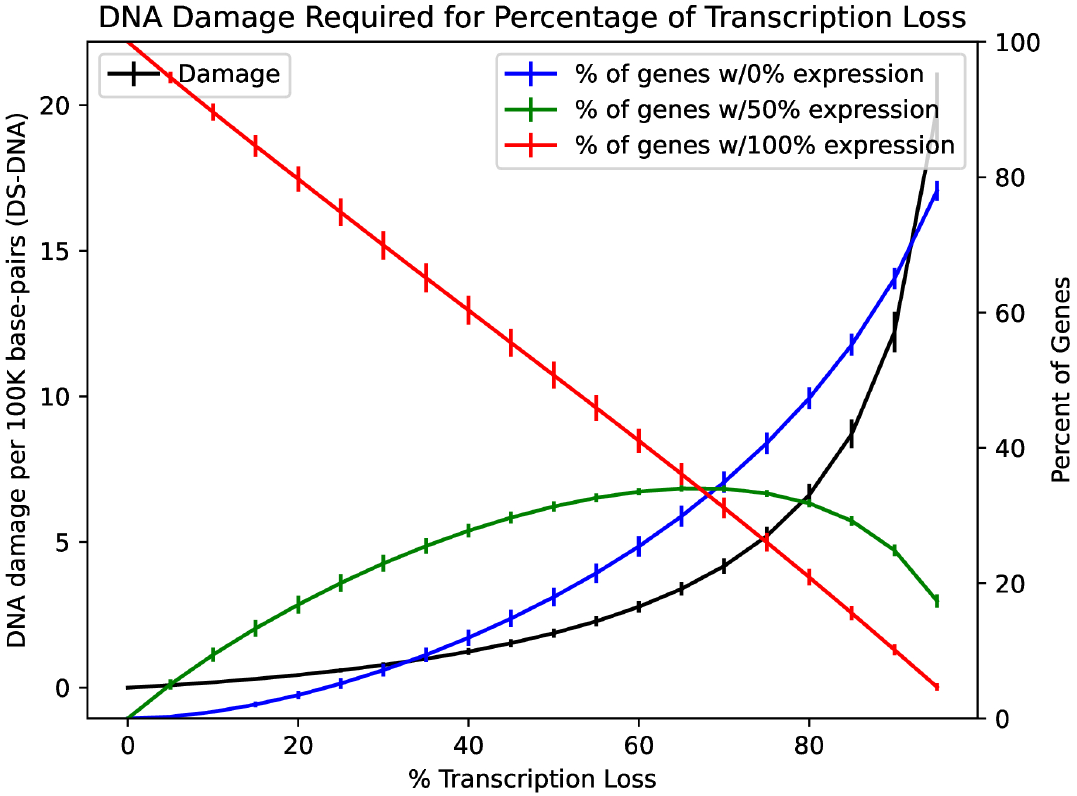
Relationship between the percentage of fully transcribed genes (blue), genes with 50 percent transcription loss (green) and genes with completely blocked transcription (red) and the number of DNA damages

**Figure 2.**
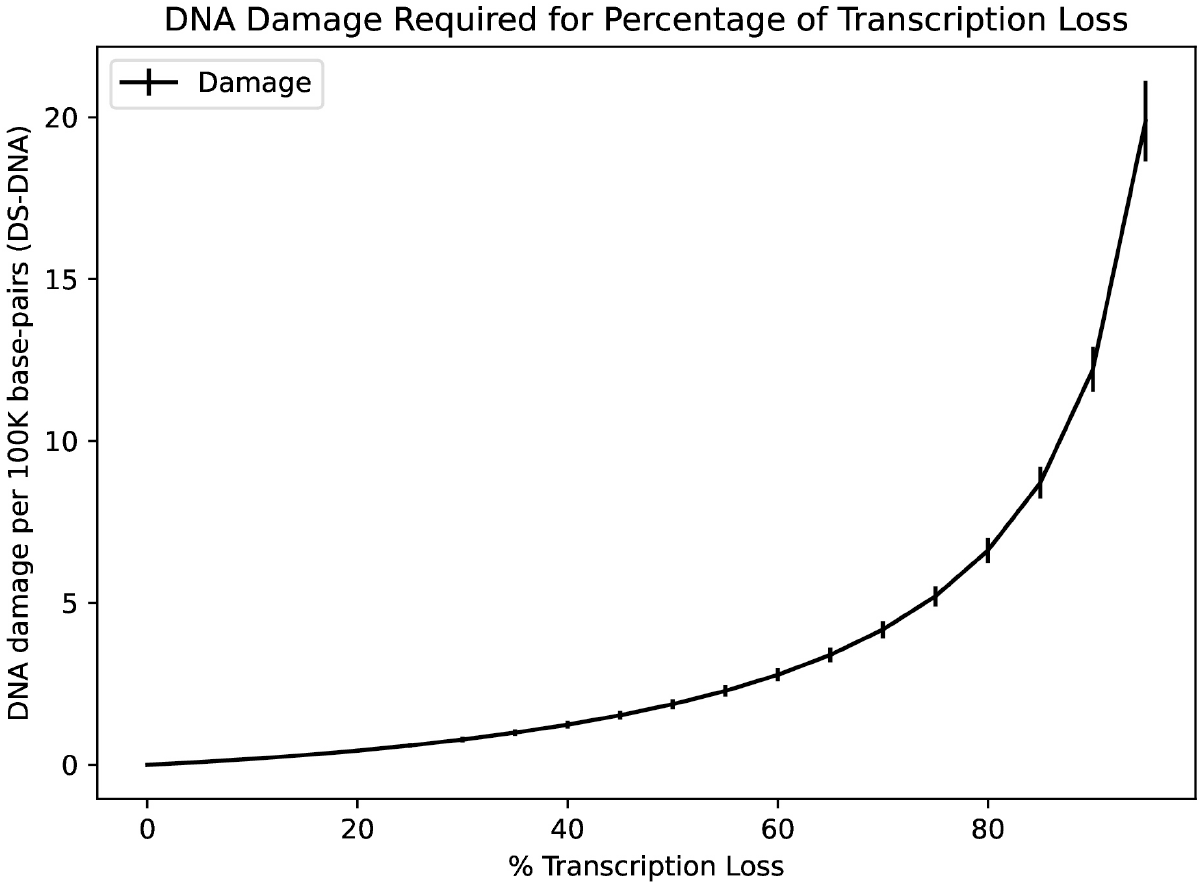
Relationship between transcription loss and a number of accumulated DNA damages

**Figure 3.**
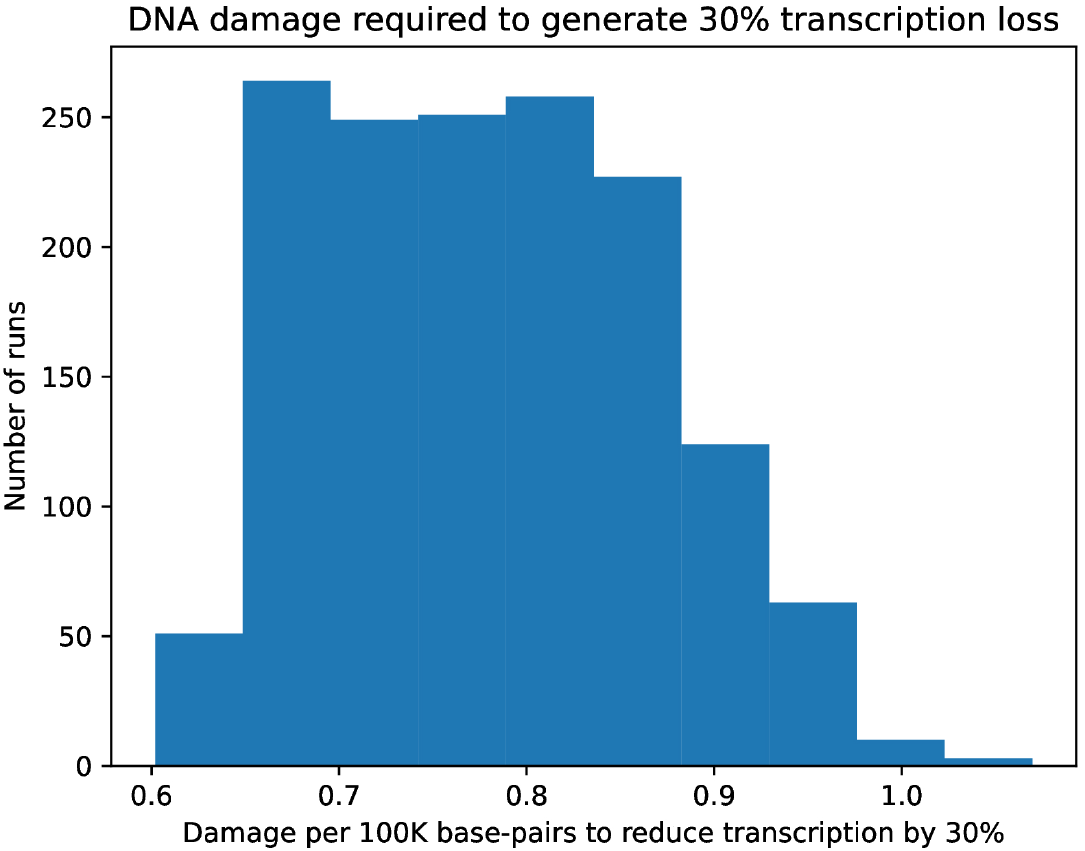
Histogram of the probability distribution of the first hitting time for the number of accumulated DNA damages resulting in 30 percent transcription loss

**Figure 4.**
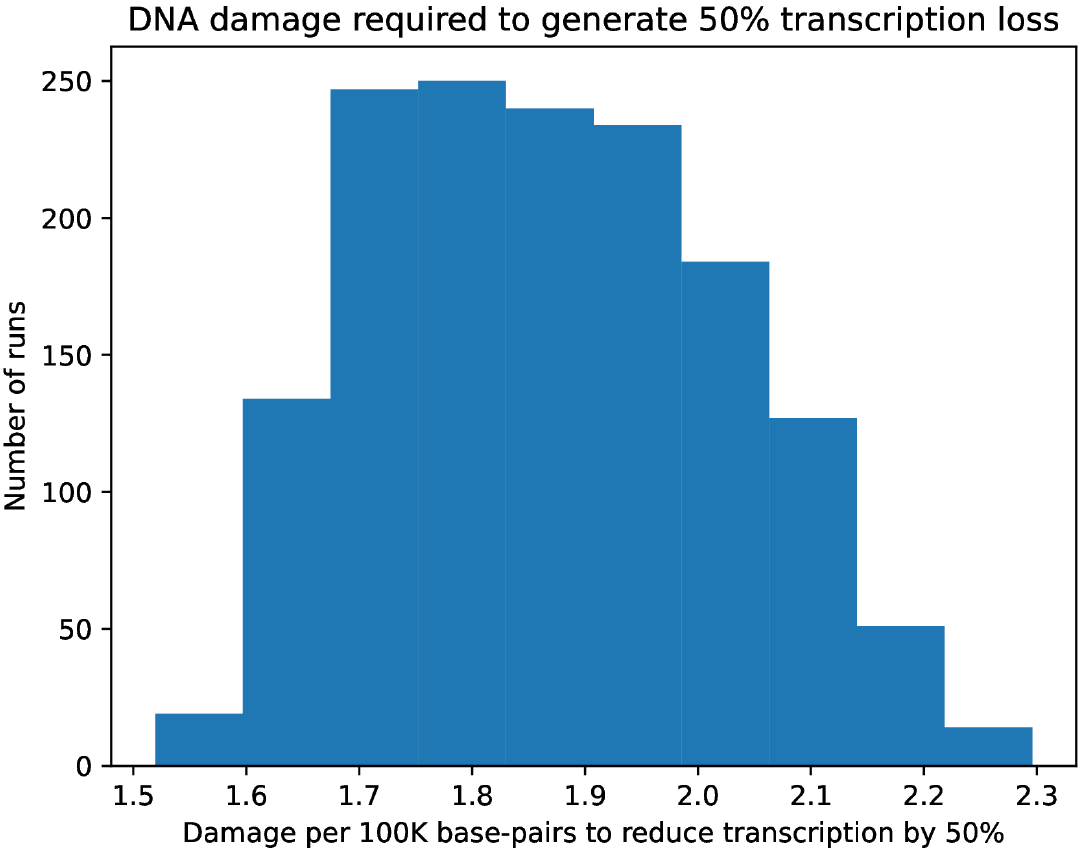
Histogram of the probability distribution of the first hitting time for the number of accumulated DNA damages resulting in 50 percent transcription loss

**Figure 5.**
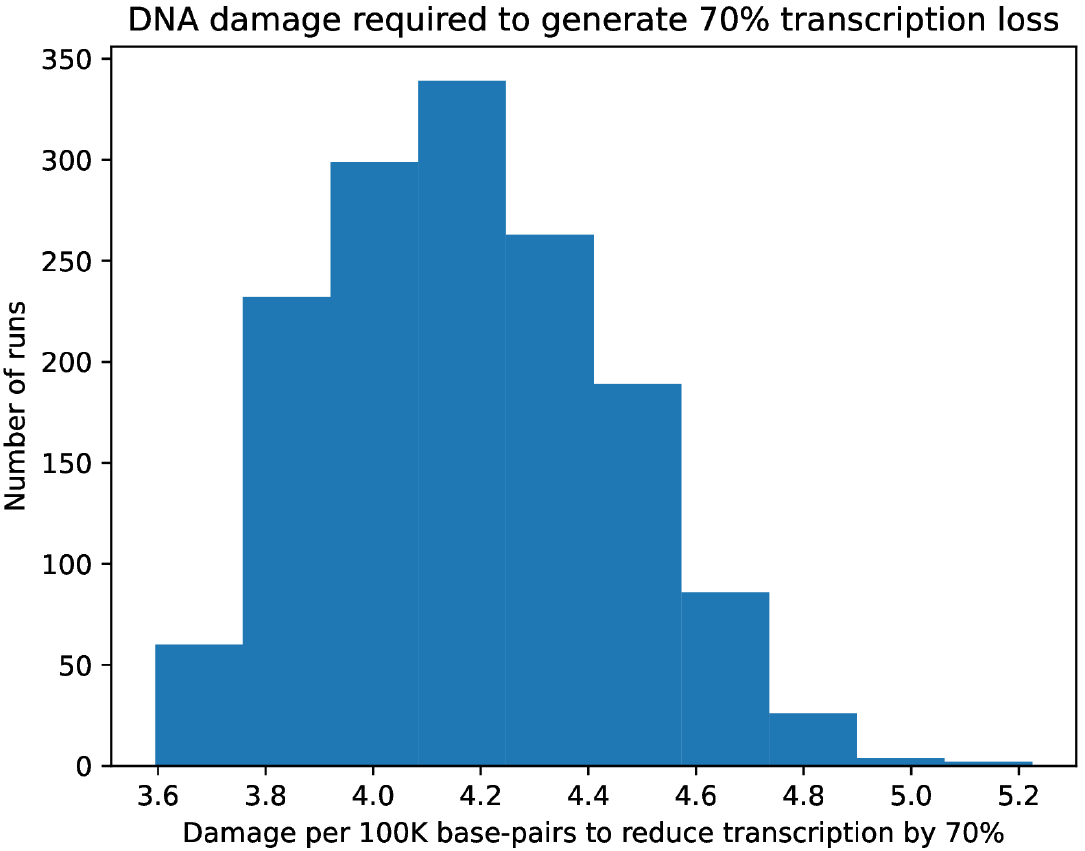
Histogram of the probability distribution of the first hitting time for the number of accumulated DNA damages resulting in 70 percent transcription loss

**Figure 6.**
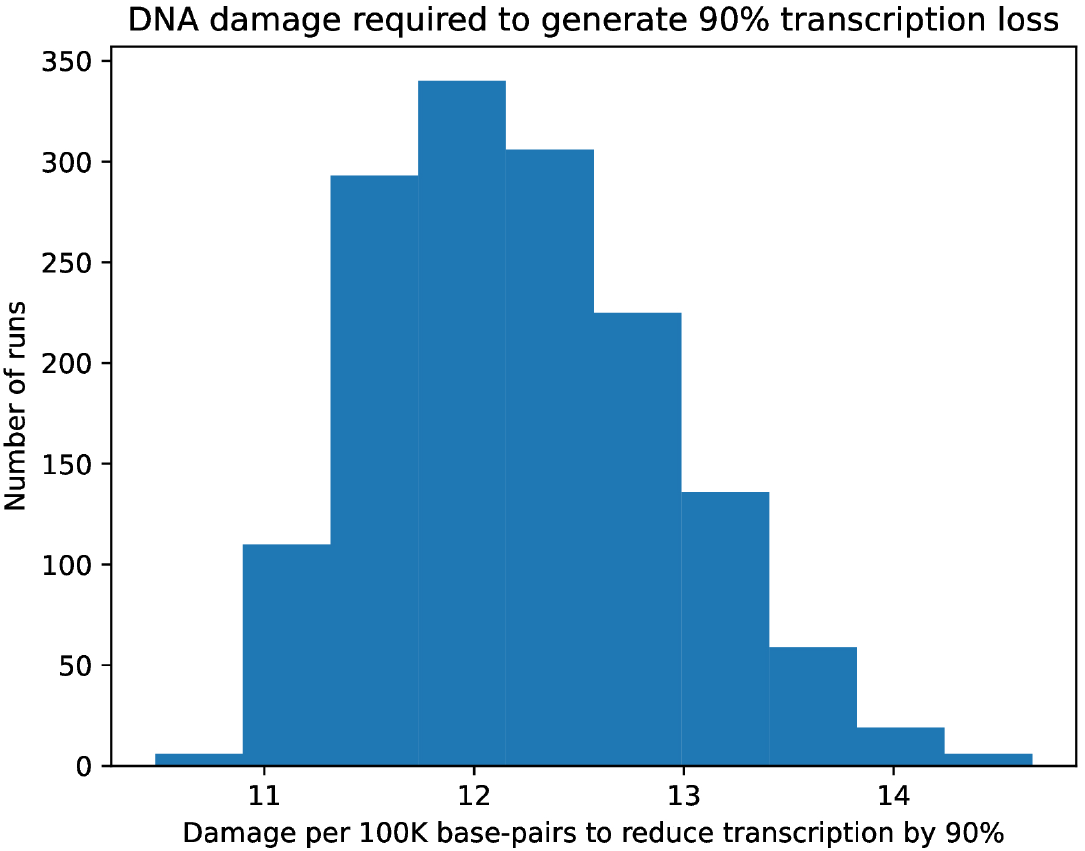
Histogram of the probability distribution of the first hitting time for the number of accumulated DNA damages needed for 90 percent transcription loss

## 5 Conclusion

We develop a simple mathematical model of DNA transcription loss due to the accumulated DNA damages. More specifically, we provide closed form formulae for the first two moments of the distribution of the transcription lost upon specified number of DNA damages. The associated stochastic process is demonstrated to converge to 1 as the number of damages tends to infinity for a variety of probabilistic convergence modes, including almost sure convergence and convergence in *L*^*p*^, for all *p* ≥ 1. Moreover, we provide closed form formulae for the probability distribution function for the random variable representing the number of damages needed to switch off both copies of a gene. Furthermore, the closed form formula for the distribution of the first hitting time of specified level of blocked transcription is provided. Unfortunately, direct application of this formula is practically infeasible due to its computational complexity, however, we have implemented a simple algorithm in a simulation study to draw statistical inference on this biologically important quantity. We plan to subsequently generalize this model further accounting for the DNA repair mechanism and study the effect on protein synthesis. Finally, we will use analytic and computational inference in conjunction with experimental in vivo and in vitro data to advance our understanding of the implications of transcription loss in aging.

